# Nanopore whole-genome sequencing reveals conserved chromosome-specific telomere architecture across tissues and populations

**DOI:** 10.64898/2025.12.10.693405

**Authors:** Niklas L. Engel, Benedikt Brors, Lars Feuerbach, Peter J. Park

## Abstract

Telomeres are repetitive nucleoprotein structures that cap the ends of linear chromosomes and are essential for maintaining genomic stability. While individual chromosome ends maintain distinct telomere lengths, the extent of this conservation across tissues and populations remains unclear due to the difficulty of analyzing repetitive telomeric sequences. Here, we show that high-coverage whole-genome Nanopore sequencing enables robust measurement of telomere length at the level of individual chromosomes. Nanopore reads yield reproducible telomere length estimates across replicates, in contrast to PacBio HiFi reads. Across > 250 individuals from 1000 Genomes and the SMaHT projects, chromosome-specific telomere length patterns are conserved across individuals and tissues, with tissues from the same individual showing highly similar patterns. This conserved landscape suggests coordinated regulation, whose disruption may contribute to genomic instability. Nanopore sequencing also allows simultaneous detection of structural variants, including disruption of *TERT* and *NHP2* that drive global telomere shortening. Furthermore, our quantification of telomere variant repeats in positional context indicates active telomerase-mediated elongation. Our integrated profiling of telomere length and structural variation enables inference of variant effects on chromosome-specific telomere dynamics and may uncover risk factors for short telomere syndromes and cancer. Importantly, positionally fully resolved telomeric variant repeat patterns may predict activated telomere maintenance mechanisms with high accuracy.

**Significance:** Resolving chromosome-specific telomere length and variant-repeat architecture across tissues and individuals provides a framework to dissect coordinated telomere maintenance, its disruption by genetic variants, and how this shapes telomere mosaicism and disease risk.

## Main

The ends of linear chromosomes are capped by repetitive DNA sequences called telomeres, which protect chromosomes from degradation and maintain genomic stability [1]. Mammalian telomeres consist of tandem repeats of the canonical hexamer TTAGGG (t-type) [2], interspersed with telomere variant repeats (TVRs) such as TCAGGG (c-type), TGAGGG (g-type), and TTGGGG (j-type), along with their reverse complements, which occur more frequently near subtelomeric regions [3]. Telomeres progressively shorten with each cell division, making telomere length a biomarker of cellular aging [4]. When telomeres fall below a critical threshold known as the Hayflick limit [5], they trigger growth arrest known as replicative senescence. If this checkpoint is bypassed, further telomere erosion leads to telomere uncapping, which activates a DNA damage response similar to that induced by double-strand breaks [6]. To prevent telomeres from shortening beyond the critical limit, certain cell types such as germ cells, stem cells, and most cancer cells activate telomerase, an enzyme that elongates and maintains telomere length [7,8]. Telomerase is a ribonucleoprotein reverse transcriptase composed of a catalytic subunit, TERT, and an RNA component, TERC, which serves as the template for repeat synthesis [9].

Telomere length is not uniform across the genome; instead, it exhibits chromosome end-specific variation. Early studies in mice and humans revealed that certain chromosome arms consistently harbor longer or shorter telomeres, indicating chromosome-specific regulation of telomere maintenance [10,11]. These length patterns are remarkably stable across tissues—including fibroblasts, peripheral blood lymphocytes, and bone marrow—within individuals, and show only limited variation between individuals [11]. Chromosome-specific telomere patterns persist through many cell divisions and appear to be established early in life. Differences can exceed 6 kilobases between chromosome ends, with conserved length distributions evident even in newborns [12]. Functionally, telomere length at individual chromosome ends can have critical consequences: it is the shortest telomere—not the average—that triggers replicative senescence and chromosomal instability [13,14]. In telomerase-deficient mice, critically short telomeres promote genome instability and premature aging, effects that can be reversed by telomerase restoration [15]. Telomere length variation across chromosome ends also carries clinical relevance. Short telomeres at Xp and 15p have been linked to increased breast cancer risk in premenopausal women [16], while unusually long telomeres on specific arms have been proposed as clonal markers in leukemia [17].

Studying telomere length at individual chromosome ends is challenging due to the highly repetitive nature of telomeres. Experimental methods are often labor-intensive, low-throughput, or restricted to a few chromosome arms, while conventional computational approaches using short-read whole-genome sequencing (WGS) provided only genome-wide averages by counting canonical 6-mer motifs rather than chromosome-specific measurements [18]. Such bulk estimates have been applied to study population variation and heritability, genome-wide associations with genetic variants, and differences between tumors and matched normal tissues [19–21]. Advances in long-read sequencing technology, however, have enabled the generation of reads with lengths of several tens to hundreds of kilobases [22]. These long reads can potentially capture the full telomeric region along with sufficient subtelomeric sequence to uniquely assign each telomere to its corresponding chromosome arm [23]. Although targeted protocols to enrich for telomeric reads have been developed for both the PacBio HiFi [24] and Oxford Nanopore Technologies (ONT) [12,23,25] platforms, such approaches come at the cost of losing genome-wide sequence information.

Here, we utilize long-read WGS to characterize chromosome-specific telomere length patterns in > 200 human individuals from diverse ancestries—the largest cohort to date for such an analysis—as well as across several endoderm- and ectoderm-derived tissues. We benchmark the accuracy and reproducibility of Telogator2 [26], a method that clusters telomeric reads based on their TVRs to assign them to specific chromosome arms, across both the PacBio HiFi and ONT platforms. We demonstrate how combined analysis of telomere length and structural variants (SVs) using Nanopore data can reveal the impact of SVs on telomere length, including the identification of likely telomere dysfunction in individual genomes. We also explore the predictive potential of TVR profiles for inferring telomere elongation dynamics and telomere maintenance mechanisms (TMMs) in cancer.

## Results

### Overview of data and analysis workflow

Our analysis utilizes three long-read datasets: WGS data generated by the 1000 Genomes project (1KG) and the Somatic Mosaicism across Human Tissues (SMaHT) project and targeted sequencing of telomeres by Schmidt *et al.* (2024). The 1KG dataset comprises ONT sequencing for 250 individuals of diverse ancestries [27], providing population-scale analysis of chromosome-specific telomere lengths across individuals and the role of SVs in genes associated with telomere regulation. The SMaHT Network aims to profile somatic mosaicism across multiple tissues from 150 healthy human donors using multiple sequencing technologies [28]. As part of its benchmarking effort, long-read data were generated for four donors. Two tissue homogenates were sequenced for two donors (liver and lung, and lung and colon), and one tissue homogenate (brain) was sequenced for the other two donors. Each sample was sequenced using both ONT and PacBio and, for each platform, at two sequencing centers, resulting in two ONT datasets and two PacBio HiFi datasets per sample. This design provides a unique opportunity to assess the reproducibility of long-read sequencing across platforms and sequencing centers. The Schmidt *et al.* (2024) dataset was generated by applying an ONT-based telomere enrichment protocol [25] to multiple human cell types, notably five TERT+ and five ALT+ cancer cell lines, as well as fibroblasts and iPSCs from six donors. The high telomeric coverage makes these data particularly suitable for positional profiling of TVRs along chromosome ends. An overview of datasets and their uses is found in Fig. 1a.

**Figure 1.**
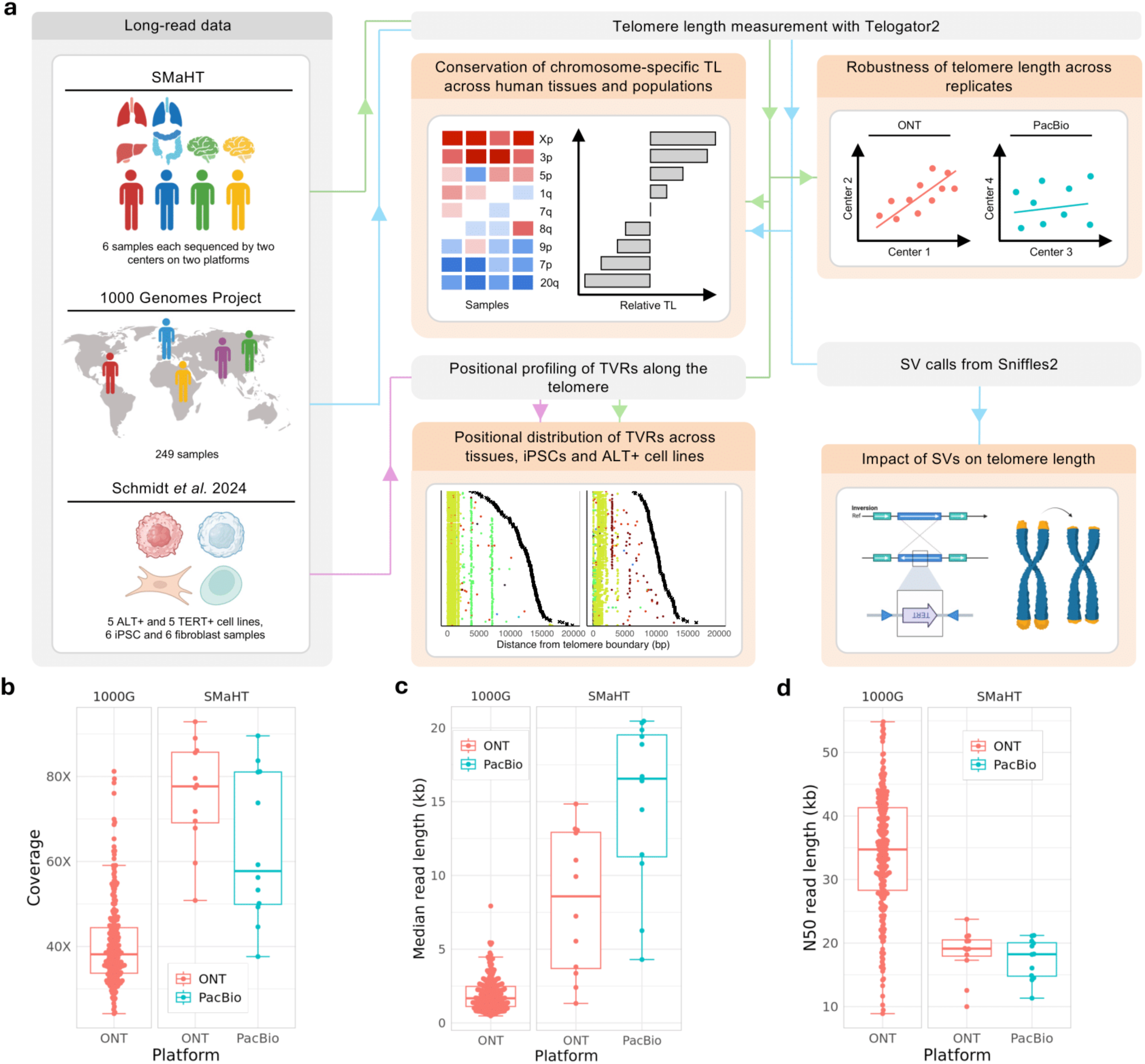
Overview of data and analysis workflow. **(a)** We calculate chromosome-specific telomere lengths for each sample in the 1KG and SMaHT datasets. We measure reproducibility across centers and platforms using the SMaHT samples. We investigate the locations of telomere variant repeats (TVRs) using the SMaHT and targeted Schmidt et al. (2024) data, and the impact of structural variants (SVs) using the 1KG data. **(b–d)** Distribution of sample properties for the datasets analyzed, including ONT data from the 1000 Genomes Project and ONT and PacBio HiFi data from SMaHT. Reported are: (b) coverage, (c) median read length, and (d) N50 read length.

As the two long-read WGS datasets were generated by different studies and sequencing centers, their properties varied. The 1KG ONT data had an average coverage of 40.5x, whereas the SMaHT ONT and PacBio datasets showed higher average coverages of 75.7x and 63.3x, respectively (Fig. 1b). Coverage varied considerably across samples, but the SMaHT datasets generally had higher coverage, reflecting their target range of 60-100x. Read length distributions also differed (Fig. 1c). The 1KG ONT samples were more homogeneous, with a median read length of 1.9 kb. By contrast, the SMaHT ONT and PacBio samples had longer, and more variable read lengths, averaging 8.2 kb and 14.9 kb, respectively. Although the short 1KG ONT read lengths and longer PacBio reads may seem unexpected, these trends are clarified by the N50 values (Fig. 1d). The 1KG ONT data had the highest N50 (34.2 kb), compared to 18.5 kb for SMaHT ONT and 17.5 kb for SMaHT PacBio. These differences highlight distinct read length distributions. For SMaHT PacBio, the median and N50 were similar, indicating tightly clustered read lengths. In contrast, ONT data contained many short reads, lowering the median, alongside very long reads that raised the N50. Of note, the target read lengths for SMaHT ONT ranged from 15-27.5 kb, and for PacBio from 17.5-20 kb. As accurate telomere length estimation requires many sufficiently long reads to cover different parts of the repeats, these properties are critical for downstream analyses.

### Robust estimation of chromosome-specific telomere length from ONT long-read WGS

Measuring telomere length from sequencing data is inherently difficult due to both biological variation and technical limitations. Telomere length can vary by several kilobases even among single cells of the same type within an individual, resulting in substantial measurement heterogeneity in bulk samples [29]. Although long-read sequencing technologies can theoretically span entire telomeric regions and extend into subtelomeric areas, it is difficult to determine with certainty that a read reaches the actual end of a telomere, since the telomere end’s 3′ G-rich single-stranded overhang is typically lost during sequencing [24].

We used Telogator2, which clusters telomeric reads based on their TVR composition to assign reads to individual chromosome arms [26]. A sufficient number of telomeric reads per chromosome end proved essential for robust estimation, and we required a minimum of 10 telomeric reads per end, which yielded a coefficient of variation below 0.15 (Fig. 2a). As expected, the number of telomeric reads correlated with the total sample coverage (Fig. 2b). The yield of telomeric reads per unit of coverage was higher for ONT (Fig. 2b), which means that the ONT samples contained more telomeric reads per chromosome end. Although the read length is similar between the two platforms (Fig. 1c,d), ONT reads captured greater telomeric content (Fig. 2c), with ONT reads extending further into telomeric regions (Fig. 2d). We note that recent improvement in ONT chemistry has also positively impacted telomere analysis. Whereas older ONT R9 chemistry exhibited basecalling artifacts in telomeric regions [30], our comparison of the abundance of canonical telomeric motifs and known error-prone motifs (TTAAAA, CTTCTT, CCCTGG and their reverse complements) between R9 and R10 ONT data for the same sample confirmed that the level of artifacts in the latest chemistry is minimal (Supplementary Fig. 1).

**Figure 2.**
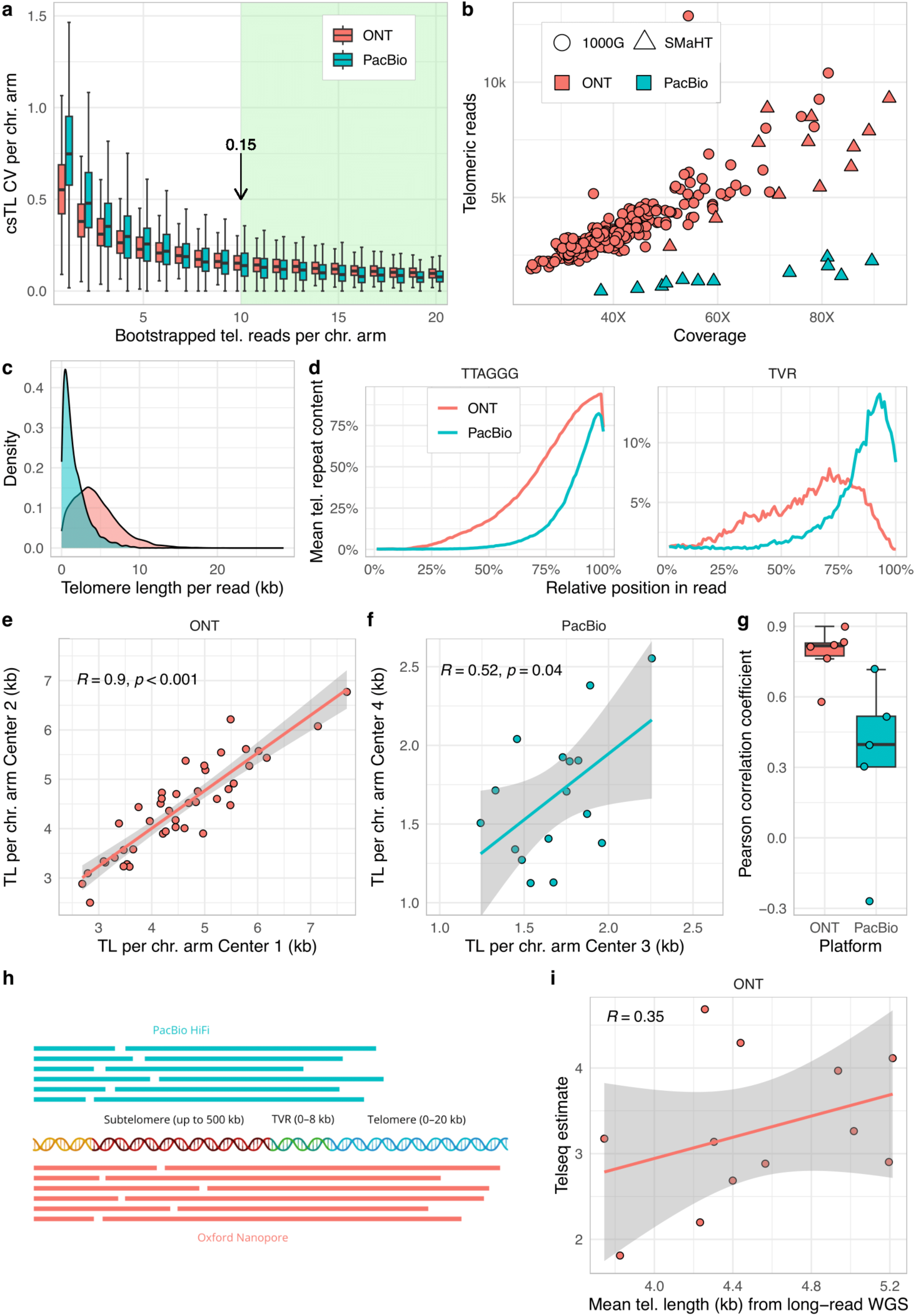
Benchmarking chromosome-specific telomere length estimation robustness from ONT and PacBio HiFi. **(a)** Boxplots showing the coefficient of variation (CV) in telomere length per chromosome arm (csTL) with increasing numbers of bootstrapped telomeric reads, separately for ONT and PacBio. **(b)** Coverage versus number of telomeric reads per sample. Dataset origin (SMaHT or 1000 Genomes Project) is indicated by point shape, and sequencing platform (ONT or PacBio) by color. **(c)** Density plot from SMaHT data showing ONT and PacBio telomeric reads by estimated telomere length per read. **(d)** Distribution of sequence composition along telomeric reads, shown as the percentage of each of 100 equal-length bins per read consisting of t-type repeats or TVRs for ONT and PacBio. **(e–f)** Correlation of telomere length per chromosome arm in SMaHT sample ST001–1A between Center 1 and Center 2 for (e) ONT and (f) PacBio data. **(g)** Pearson correlation coefficients of chromosome-specific telomere length estimates for a set of matched samples sequenced at two different centers for ONT and PacBio. **(h)** Schematic illustrating read mapping across subtelomeric regions, TVRs, and the canonical telomere, comparing ONT and PacBio read coverage. **(i)** Correlation of mean telomere length estimates from ONT with matched short-read WGS estimates using Telseq. Regression lines and Pearson correlation coefficients are shown.

To assess the robustness and reproducibility of these estimates across data generators and platforms, we analyzed identical biological samples that were independently sequenced at different centers using the same platform (either ONT or PacBio HiFi). For a SMaHT sample (ST001-1A), ONT estimates correlated strongly between replicates, whereas PacBio estimates did not (Fig. 2e,f). Across all SMaHT benchmarking samples, ONT consistently showed higher inter-replicate correlations than PacBio (Fig. 2g). Perhaps not surprising given the lack of consistent estimates on PacBio, correlations between ONT- and PacBio-derived estimates from the same sample were generally low (Supplementary Fig. 2). These results indicate that ONT achieves better coverage of telomere content and thus provides more accurate estimates (Fig. 2h). Our subsequent analysis is therefore base only on ONT data.

The long-read data we have enable us to assess whether the sample-level telomere length estimates from Illumina short-reads are reliable—as short reads cannot be mapped to specific chromosomes, short-read estimates provide a sample level essentially by counting the number of the TTAGGG (or CCCTAA) motif. Across the 4 SMaHT tissues, we find that short-reads (using TelSeq [31]) estimates have only a weak positive correlation (r=0.35) with ONT data (Fig. 2i; the correlation with PacBio is negative (Supplementary Fig. 3)). Although short read-based estimates have been used in multiple studies [19–21], our analysis suggests that only general trends might be discernable from the data.

### Chromosome-specific telomere length is conserved across human tissues

Having established that whole-genome ONT data enable robust estimation of chromosome-specific telomere lengths, we next examined their variation across chromosome arms in the six SMaHT samples, representing four donors of different ages and tissues. Across tissues, we observed clear differences in mean telomere length, with lung samples from donors ST001 and ST002 showing the longest mean telomeres, while brain samples from donors ST003 and ST004 had the shortest. Although the sample size here is too small to draw any conclusion, we did not observe a strong negative correlation between age and mean telomere length as previously described [4]. For example, colon and lung from the 74-year-old donor (ST002) had longer mean telomeres than liver from the 22-year-old donor (ST001; Fig. 3a). However, when we examined the 25th percentile of telomere lengths per chromosome arm, an age-related pattern emerged: younger donors (ST001, ST003) had consistently longer telomeres than older donors (ST002, ST004; Fig. 3b). This discrepancy between mean and 25th percentile values suggested that in the underlying distribution of single reads, older individuals had a larger fraction of short telomere reads compared the younger individuals (Fig. 3c) and that the 25th percentile may better capture age-associated telomere attrition.

**Figure 3.**
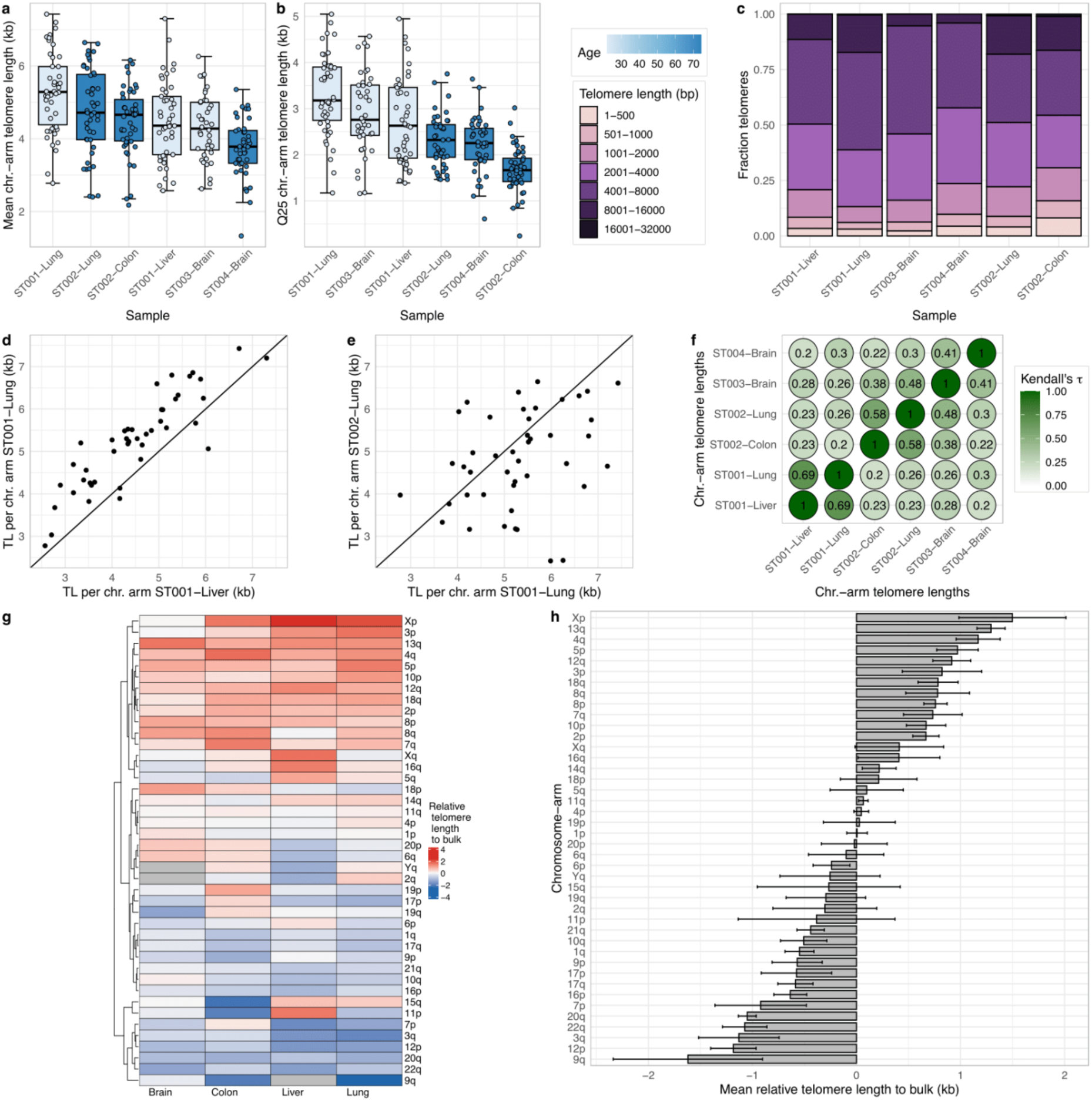
Distribution of chromosome-specific telomere lengths across human tissues. **(a–b)** Telomere length distributions per chromosome arm across six samples derived from liver, lung, colon, and brain tissues of donors ST001–ST004, colored by donor age. Telomere lengths are summarized by the (a) mean and (b) 25th percentile. **(c)** Distribution of binned telomere lengths from telomeric reads across the six SMaHT samples. **(d–e)** Correlation of telomere length per chromosome arm in SMaHT samples (d) ST001–Liver and ST001–Lung and (e) ST001–Lung and ST002–Lung. **(f)** Kendall correlation coefficients of chromosome-specific mean telomere length estimates between all six samples. **(g)** Heatmap showing telomere lengths relative to bulk telomere length estimates per chromosome arm (kb) for brain, colon, liver, and lung. **(h)** Bar plot of mean relative to bulk telomere length per chromosome arm (kb) across the four tissues. Error bars represent the standard deviation.

We asked whether the relative ranking of chromosome arms in terms of telomere lengths is conserved across tissues, as previously reported for blood-derived samples [12]. To assess how consistently chromosome arm lengths are ordered between samples, we calculated Kendall rank correlation, a non-parametric test based on the number of concordant and discordant pairs (e.g., a pair is concordant if Sample A chr 5 > Sample A chr 7 and Sample B chr 5 > Sample B chr 7). The strongest correlations occurred between tissues from the same individual—liver and lung in ST001 (Kendall’s τ = 0.69; Fig. 3d), and colon and lung in ST002 (Kendall’s τ = 0.58)—while the same tissue between different individuals were not well correlated (ST001 and ST002 lungs; Kendall’s τ = 0.26; Fig. 3e). The correlation matrix between every pair is shown in Fig. 3f.

Notably, a consistent ranking for chromosome arms emerged across different tissues. Certain arms, such as Xp, 13q, and 4q, showed consistently long telomeres, whereas others, such as 9q, 12p, and 3q, had consistently short telomeres (Fig. 3g,h). This result demonstrates that, despite the limited size of our cohort, the relative ordering of telomere lengths among chromosome arms is conserved not only in blood, as previously reported, but also across brain, colon, liver, and lung. The specific chromosomes show strong correspondence with those reported by Karimian *et al.* (2024), who found 3p, 12q, and 4q among the longest and 17p, 20q, and 12p among the shortest arms in blood. Their study did not include Xp, which we found to be among the longest.

### Chromosome-specific telomere length is conserved across human populations

We next extended the analysis of conserved telomere length ranking to a high-coverage ONT dataset from over 200 individuals in the 1000 Genomes Project [27], representing a broad range of global populations. To our knowledge, this is the largest and most ethnically diverse cohort analyzed for chromosome arm–specific telomere length.

Across individuals from five major regions—Africa, America, East Asia, Europe, and South Asia—telomere length distributions were broadly similar, though European individuals showed notably shorter telomeres, with median lengths nearly 50% lower than in other populations (Fig. 4a). Stratification by sex revealed consistently longer telomeres in females than in males across all regions except Europe, where no significant difference was detected (Fig. 4b). At the individual level, telomere lengths varied widely, ranging from about 1 kb to over 15 kb (Fig. 4c), underscoring the substantial inter-individual diversity in blood telomere length.

**Figure 4.**
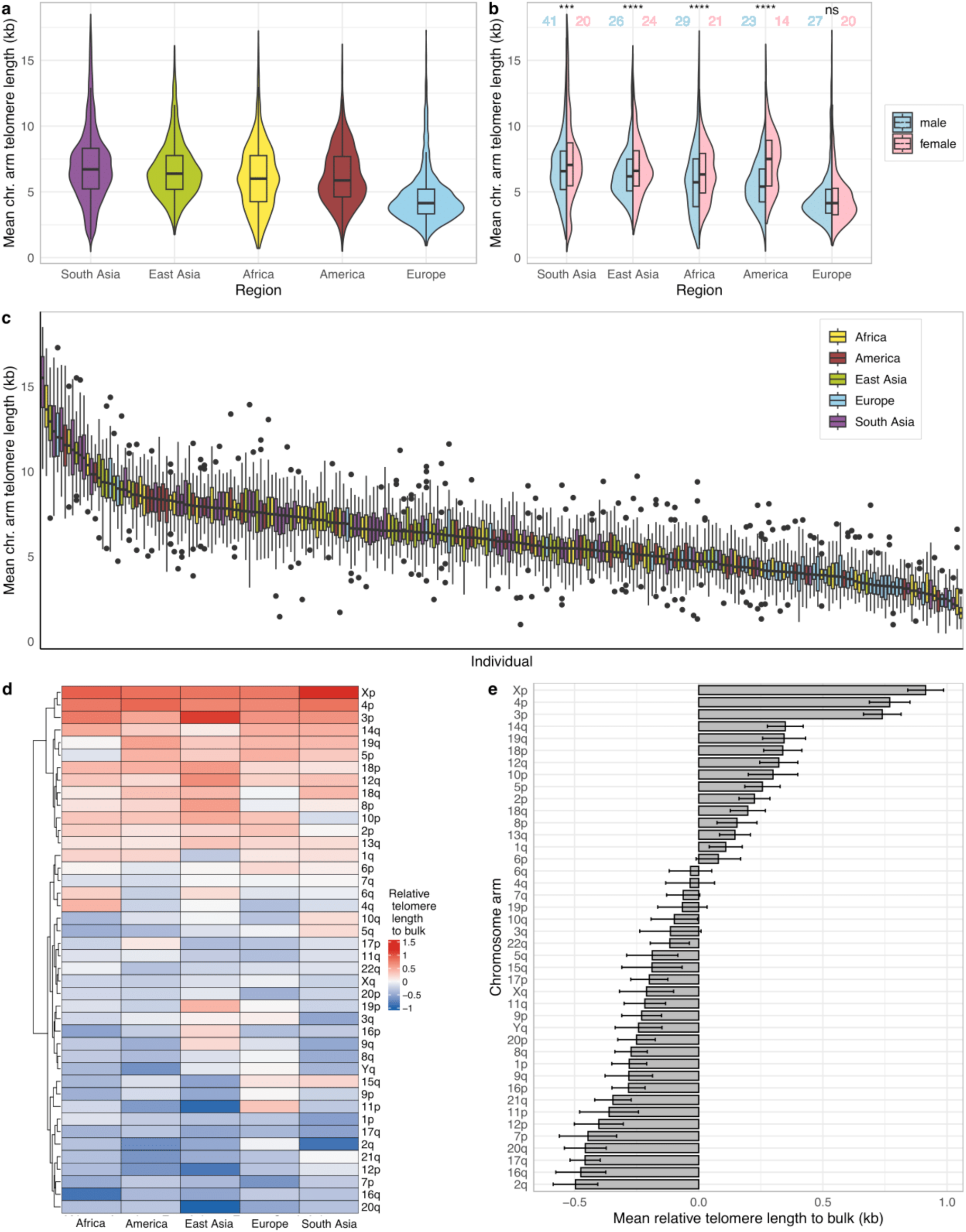
Distribution of chromosome-specific telomere lengths across human populations. **(a)** Distribution of mean telomere length per chromosome arm across individuals from Africa, America, East Asia, Europe, and South Asia. **(b)** Same as (a), stratified by sex (male: blue; female: pink). P-values from Wilcoxon rank-sum tests comparing sexes are indicated above each boxplot. **(c)** Per-individual distribution of mean telomere length per chromosome arm, colored by region and ranked in descending order. **(d)** Heatmap of telomere lengths relative to bulk estimates per chromosome arm, aggregated across individuals within each region. **(e)** Mean relative telomere length per chromosome arm across individuals from the five regions. Error bars indicate the standard deviation.

We next examined whether telomere length ranking among chromosome arms was conserved across population groups. Similar to the tissue-level results, certain arms remained consistently long or short across all five regions: Xp, 3p, and 4p showed the longest telomeres, whereas 12p, 2q, and 20q were consistently among the shortest (Fig. 4d,e). This conserved rank order moderately aligned with that observed across the four primary tissue types, suggesting that chromosome-arm-specific telomere length is a stable genomic feature across both individuals and populations. Minor discrepancies likely reflect the limited sample size of the tissue dataset. Overall, these findings indicate that telomere length variation at individual chromosome arms is a conserved property of the human genome across diverse populations and tissues representing all three germ layers.

### Large inversions spanning *TERT* and *NHP2* shorten telomere length across all chromosome ends

A key advantage of long-read WGS over telomere enrichment-based methods is the preservation of broader genomic context. We leveraged SV calls from the high-coverage ONT 1000 Genomes Project cohort, for which we have estimated chromosome-specific telomere lengths, to test whether SVs overlapping known telomere-regulating genes were associated with altered telomere length. This joint analysis within the same individuals enabled us to investigate potential genetic determinants of telomere length variability.

Among more than 200 individuals, we identified two SVs with clear relevance to telomere biology. One individual (HG03166) carried a large inversion spanning *TERT*, which encodes the catalytic reverse transcriptase subunit of telomerase (Fig. 5a). Another (HG00237) harbored an inversion encompassing *NHP2*, a core component of the H/ACA small nucleolar ribonucleoprotein complex essential for telomerase RNA stability (Fig. 5b). Both inversions—the two outliers in terms of the magnitude and statistical significance of decreased telomere lengths (Fig. 5c)—are likely to disrupt telomerase function and compromise telomere maintenance. Consistent with this expectation, these individuals exhibited the shortest and second shortest mean telomere lengths in the entire cohort (Fig. 5d). Compared with population-matched controls, the individual with the *TERT* inversion showed substantially reduced telomere lengths across all chromosome arms, as shown by the distribution of lengths from individual telomeric reads (Fig. 5e,f). A similar pattern was observed for the *NHP2* inversion carrier (Fig. 5g,h). Together, these findings demonstrate that SVs disrupting key telomere regulators can cause pronounced, genome-wide telomere shortening.

**Figure 5.**
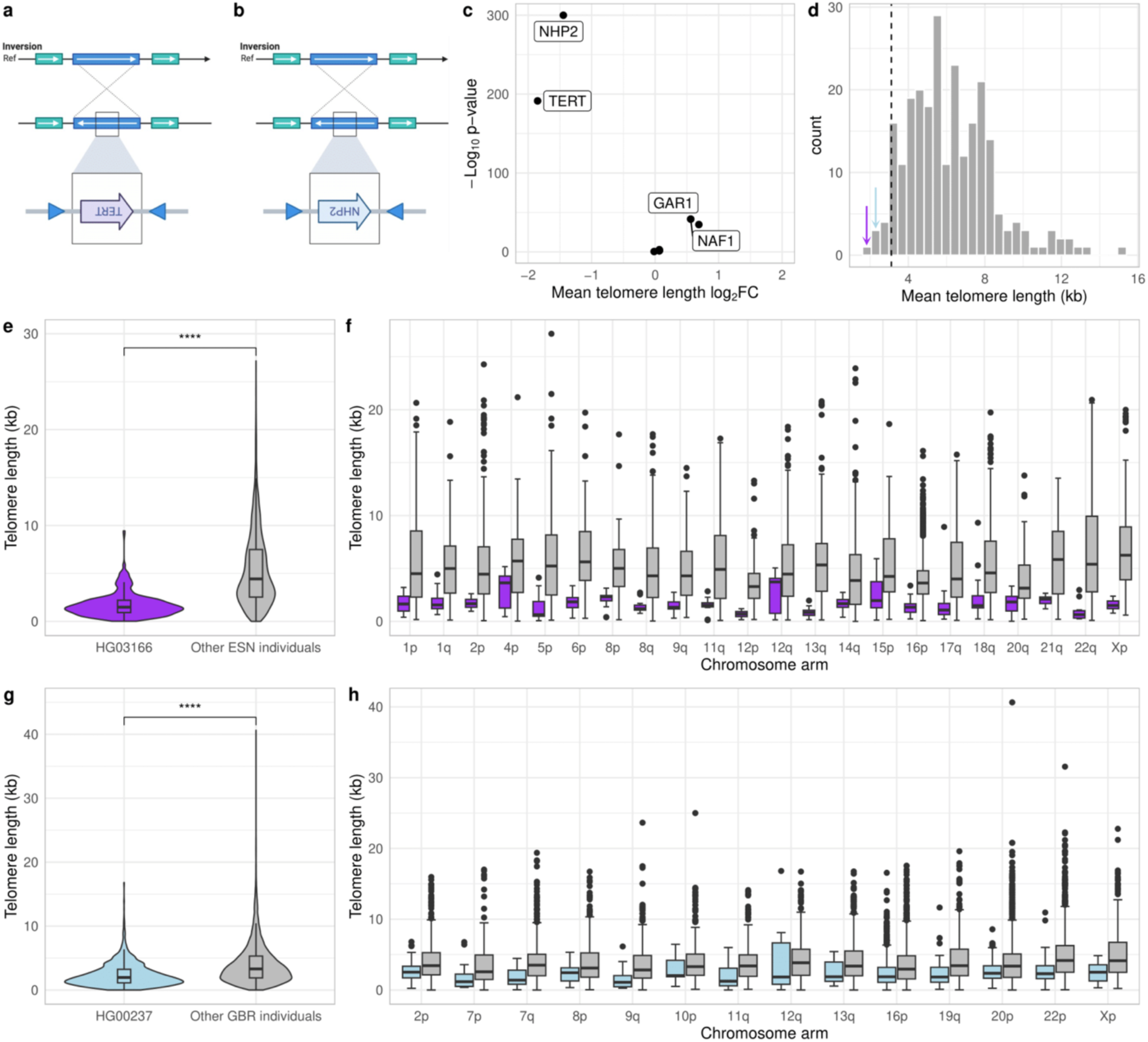
Telomere length distribution in individuals with TERT and NHP2 inversions. **(a–b)** Schematic representation of large inversion events spanning the entire genes of (a) TERT in individual HG03166 and (b) NHP2 in individual HG00237. **(c)** Fold changes in mean telomere length and associated p-values for individuals harboring structural variants overlapping promoters of selected telomere-regulation-related-genes, compared to the rest of their respective population. **(d)** Histogram of the mean telomere length per individual in the cohort. The black dashed line marks the 5th percentile; colored arrows indicate mean telomere lengths of HG00237 (light blue) and HG03166 (purple). **(e–h)** Telomere length distributions comparing individuals with the inversion to others in their population: (e, g) all telomeric reads and (f, h) telomeric reads per chromosome arm for HG03166 (Esan in Nigeria (ESN) population; e–f) and HG00237 (British from England and Scotland (GBR) population; g–h).

### Tissue-invariant telomere variant repeat (TVR) patterns and their preservation during telomere elongation

TVRs are interspersed throughout the telomere, typically enriched near the subtelomeric boundary and becoming sparser toward the distal end (Fig. 6a). They are thought to arise from replication errors during telomere extension by telomerase [32]. Although short-read sequencing can quantify overall TVR abundance, long reads uniquely provide positional resolution, allowing discrimination between subtelomeric and intratelomeric TVRs.

**Figure 6.**
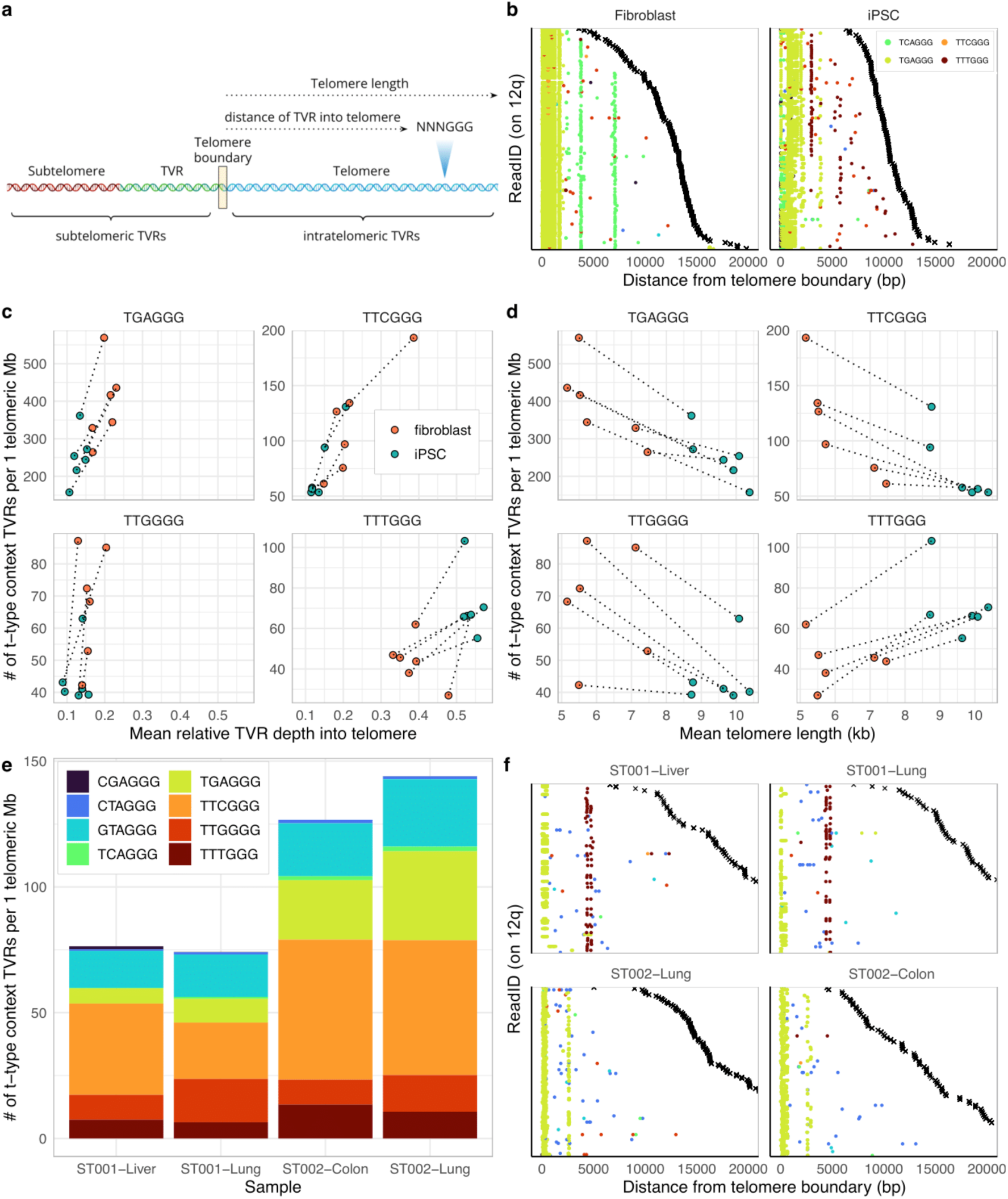
Distribution of TVRs across telomeres in human tissues and during telomere elongation captured from long reads. **(a)** Schematic of TVR localization relative to the telomere boundary in long reads. TVRs upstream of the telomere boundary, often in TVR-rich or subtelomeric regions, are classified as subtelomeric; those downstream are considered intratelomeric. Distance to the boundary is measured directly from long reads. **(b)** Positional distribution of singleton TVRs (CGAGGG, CTAGGG, GTAGGG, TCAGGG, TGAGGG, TTCGGG, TTGGGG, TTTGGG) along individual long reads at chromosome 12q, aligned by distance from the telomere boundary, in donor-matched fibroblasts and iPSCs from the 94-year-old donor. Black crosses indicate the end of each telomeric read. **(c–d)** Number of singleton TVRs per 1 Mb of telomeric sequence for TGAGGG, TTCGGG, TTGGGG, and TTTGGG, plotted against (c) mean relative distance within the telomere and (d) mean telomere length (kb) for each sample. Donor-matched fibroblast/iPSC pairs are connected by dotted lines. **(e)** Bar plot showing the number of singleton TVRs per 1 Mb of telomeric sequence for the eight selected TVRs in samples ST001-Liver, ST001-Lung, ST002-Colon, and ST002-Lung. **(f)** Positional distribution of the eight singleton TVRs in long reads at chromosome 12q for the four samples in (e), aligned by distance from the telomere boundary. Black crosses indicate the end of each telomeric read.

Here, we quantify TVRs in their positional context using long-read sequencing to profile intratelomeric variants and their dynamics in telomere elongation across human tissues. For each telomeric read, we first defined the telomere boundary as the estimated telomere length from the read end and recorded the position of every detected TVR. This enabled classification of TVRs as subtelomeric or intratelomeric and calculation of their absolute and relative distances from the boundary (Fig. 6a). Subsequent analyses focused on intratelomeric TVRs.

To study TVR dynamics during telomere elongation, we analyzed telomere-enriched ONT data from donor-matched fibroblasts and iPSCs generated by Schmidt *et al.* (2024). Aligning TVRs by distance from the telomere boundary produced pileup-like profiles of TVR occurrence along each chromosome arm (Fig. 6b). On chromosome arm 12q, fibroblast reads showed highly concordant TVR positions, with recurrent motifs appearing at consistent distances and occasional unique variants likely reflecting sequencing noise, demonstrating the positional precision of ONT-based TVR detection. Upon reprogramming to iPSCs, which reactivates telomerase, this profile shifted: the TGAGGG cluster near the boundary persisted, whereas two fibroblast-specific TCAGGG repeats disappeared. Conversely, a TTTGGG repeat appeared downstream of TGAGGG in a subset of iPSC reads. These changes indicate that telomeres shortened almost up to the TGAGGG cluster (<5 kb) and then elongated, with telomerase activity introducing new TTTGGG repeats. The partial presence of TTTGGG across reads further suggests clonal heterogeneity in telomere elongation (Fig. 6b).

Aggregating TVR occurrences across all chromosome arms reinforced these trends. TGAGGG, TTCGGG, and TTGGGG motifs decreased in abundance with elongation, while their positions remained stable or shifted slightly toward the boundary. In contrast, TTTGGG repeats increased in both abundance and distance within the telomere (Fig. 6c). Across samples, mean telomere length correlated negatively with TGAGGG, TTCGGG, and TTGGGG abundance but positively with TTTGGG (Fig. 6d). This suggests that most TVRs are not incorporated during elongation, but TTTGGG repeats likely represent *de novo* insertions by telomerase.

We next examined tissue-specific TVR profiles across tissues from the same donor in the SMaHT dataset (Fig. 6e). Donor ST002 exhibited consistently higher counts of GTAGGG, TGAGGG, and TTCGGG motifs across both tissues compared to ST001, whereas increases in TTTGGG were modest. Within individuals, absolute and relative TVR counts were similar between tissues. This pattern became even clearer at the single-telomere level (Fig. 6f). On 12q, liver and lung samples from ST001 shared nearly identical TVR profiles, including a prominent TGAGGG cluster near the boundary followed by two TTTGGG repeats. Similarly, liver and colon from ST002 displayed a TGAGGG cluster and downstream repeats in the same configuration. Some reads, however, lacked one or more downstream TVRs (Fig. 6f), which may reflect true telomeric heterogeneity arising from distinct elongation events or incomplete sequencing coverage.

The strong concordance of TVR profiles between tissues from the same donor supports a common progenitor origin, in which telomeres were elongated and retained distinct TVR patterns. The persistence of shared motifs suggests that subsequent telomere shortening has not progressed beyond the most distal shared TVR, providing a historical record of telomere elongation. Based on this, we infer a minimum historical telomere length of approximately 5 kb in ST001 (liver and lung) and approximately 3 kb in ST002 (liver and colon).

### TVR profiles predict alternative lengthening of telomeres (ALT) status at the single-molecule level

TVRs are also implicated in cancer, particularly through the ALT pathway, an alternative telomere maintenance mechanism active in approximately 4-11% of cancers [33–35]. ALT is especially prevalent in tumors such as osteosarcoma and glioblastoma and operates through recombination-based telomere extension rather than telomerase reactivation [19,34,36].

Previous studies have reported characteristic enrichment and depletion patterns of specific TVRs in ALT+ cancers, likely reflecting the mechanistic consequences of recombination-driven telomere synthesis [32,37,38]. As the ALT pathway is increasingly explored as a therapeutic target [39], reliable and scalable approaches for determining ALT status are of growing interest. To assess whether long read-derived TVR profiles could serve this purpose, we analyzed telomere enrichment data from Schmidt *et al.* (2024) comprising five ALT+ and five TERT+ cell lines. We focused on the positional distribution of TVRs within telomeres to evaluate their potential as markers of ALT activity.

We first compared the total counts and positional distance profiles of intratelomeric TVRs across all chromosomes in ALT+ and TERT+ cell lines. The TERT+ lines exhibited a pattern consistent with that seen in iPSCs—a strong increase in both the abundance and distal positioning of TTTGGG repeats relative to the telomere boundary, consistent with active telomerase-mediated elongation (Fig. 7a). For most other TVRs, there were no clear global differences between ALT+ and TERT+ lines. However, three TVRs—TCAGGG, TGAGGG, and TTCGGG—showed striking outlier enrichment in specific ALT+ cell lines, with counts exceeding those of all other lines by more than tenfold. Distinct ALT+ lines were enriched for different TVRs: Saos-2 for TCAGGG, G-292 for TGAGGG, and U2-OS for TTCGGG (Fig. 7a). Notably, overall TVR abundance did not correlate with mean telomere length across ALT+ and TERT+ samples (Supplementary Fig. 4).

**Figure 7.**
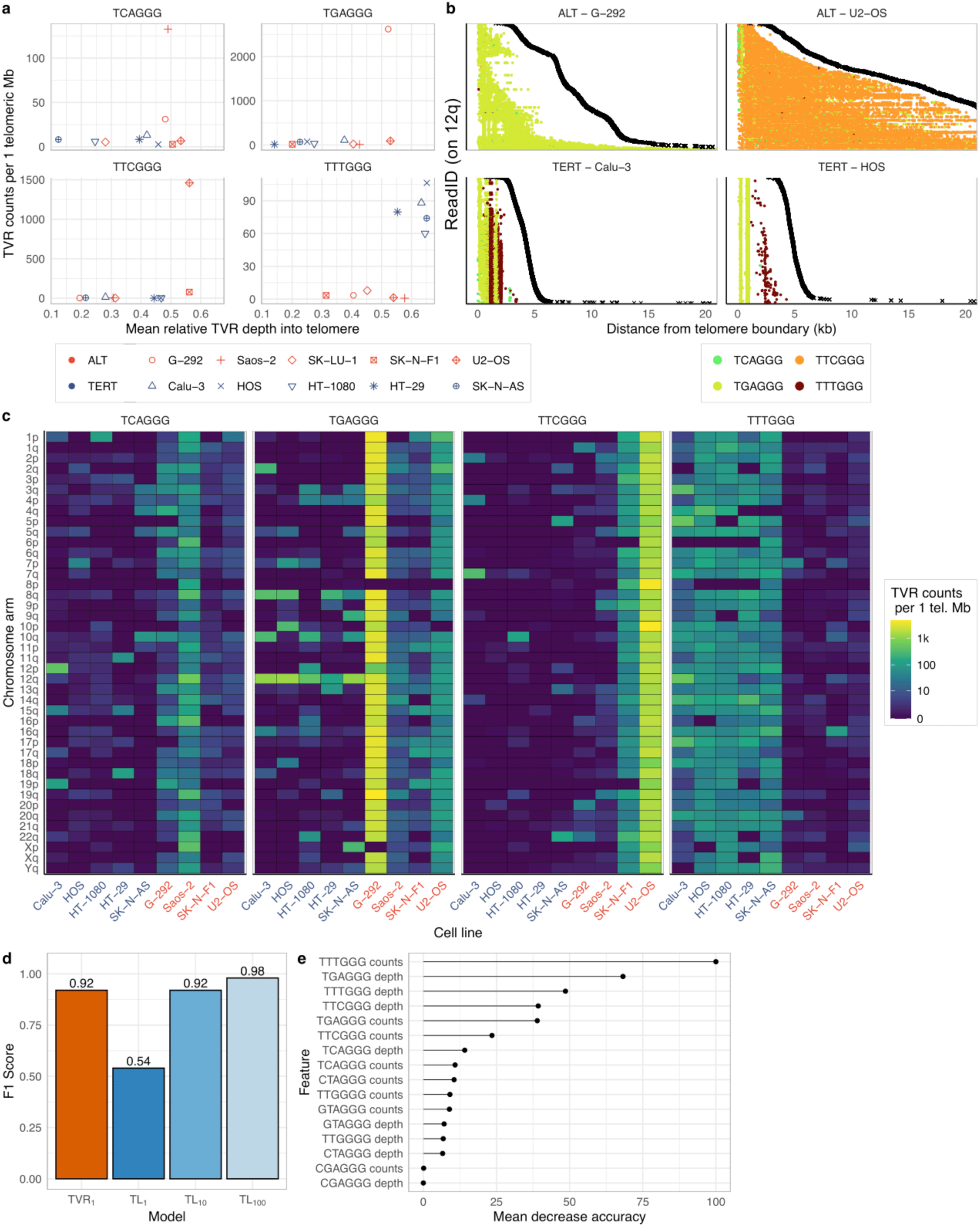
Distribution of telomere variant repeats in ALT+ vs TERT+ cell lines and prediction of ALT status. **(a)** Number of singleton TVRs (TCAGGG, TGAGGG, TTCGGG, TTTGGG) per 1 Mb of telomeric sequence plotted against the mean relative distance within the telomere per sample. ALT+ and TERT+ status is indicated by color; cell lines are distinguished by shape. **(b)** Positional distribution of the four TVRs shown in (a) along individual long reads at chromosome 12q for ALT+ cell lines (G-292, U2-OS) and TERT+ cell lines (Calu-3, HOS), aligned by distance from the telomere boundary. Black crosses indicate the end of each read. **(c)** Number of singleton TVRs per 1 Mb of telomeric sequence per chromosome arm for the four TVRs across ALT+ and TERT+ cell lines. **(d)** F1 scores from machine learning models classifying long reads as originating from ALT+ or TERT+ cell lines. The TVR_1_ model is a random forest trained on normalized counts and telomeric distance of eight TVRs from individual long reads. TL_1_, TL_10_, and TL_100_ are generalized linear models using telomere length alone (TL_1_), or combined with mean and coefficient of variation of telomere length from sets of 10 (TL_10_) or 100 (TL_100_) reads. **(e)** Feature importance in the TVR_1_ model, shown as mean decrease in accuracy, for count and positional distance features of eight selected TVRs.

We next asked whether the enrichment of specific TVRs in ALT+ lines was restricted to individual loci or distributed genome-wide. To resolve these patterns at higher resolution, we examined the TVR composition of chromosome arm 12q in two ALT+ and two TERT+ lines (Fig. 7b). As expected, TERT+ lines displayed the canonical elongation-associated pattern observed in iPSCs. In contrast, the ALT+ lines showed highly distinctive and pervasive TVR signatures: G-292 displayed widespread TGAGGG enrichment, while U2-OS exhibited pervasive TTCGGG incorporation, spanning the entire telomere. Expanding the analysis genome-wide confirmed that these enrichments were not confined to specific loci: ALT+ cell lines with outlier TVR profiles displayed high abundance of their respective repeats across nearly all chromosome arms, whereas TERT+ lines showed minimal or no enrichment. Even ALT+ lines without extreme outlier status exhibited moderately elevated levels of these repeats relative to TERT+ lines. In contrast, the telomerase-associated repeat TTTGGG was consistently enriched across nearly all chromosome arms of TERT+ lines and depleted in ALT+ lines (Fig. 7c). These results demonstrate that the ALT mechanism induces pervasive, cell line–specific alterations in telomere composition, characterized by incorporation of distinct noncanonical repeats.

Finally, to evaluate the predictive potential of TVR profiles from long reads, we trained four machine learning models to classify individual reads (or sets of reads) as originating from ALT+ or TERT+ cell lines. The first model (TVR_1_) was a random forest classifier using the counts and positional information of the eight most prominent TVRs (CGAGGG, CTAGGG, GTAGGG, TCAGGG, TGAGGG, TTCGGG, TTGGGG, and TTTGGG) from a single read. The remaining three models (TL_1_, TL_10_, and TL_100_) were generalized linear models predicting ALT status from telomere length alone, using one, ten, or one hundred reads, respectively. Remarkably, the TVR_1_ model achieved high predictive performance using a single long-read, substantially outperforming the single-read length-based model (TL_1_). Comparable accuracy was only reached when length-based models were provided with 10–100 reads (Fig. 7d). Feature importance analysis revealed that both the counts and positional information of TTTGGG, TGAGGG, and TTCGGG repeats were key predictors, with TTTGGG repeat counts being most influential (Fig. 7e). Together, these findings highlight the strong diagnostic potential of TVR profiles as single-molecule biomarkers of ALT activity, offering a scalable and interpretable read-level approach for telomere-based tumor classification.

## Discussion

Telomere biology is central to understanding cellular aging and genome integrity, yet a comprehensive view of chromosome-specific telomere length and sequence variation across tissues, ancestries, and cancer contexts using long-read WGS has been lacking. Prior studies have demonstrated the utility of long-read data for telomere analysis, but several issues have not been explored, such as differences between sequencing platforms, the positional structure of TVRs at single-base resolution, and the biological conservation of telomere patterns across chromosomes.

Here, we leveraged high-coverage long-read WGS to profile telomere features across diverse human samples. We first compared sequencing platforms and found that ONT produces more reproducible chromosome-specific telomere length estimates than PacBio HiFi, although both platforms continue to improve rapidly and the results may change in the future. Unlike targeted enrichment assays, ONT WGS at 30X provides sufficient read depth for reliable estimation of telomere length and genomic variants, enabling integrated analysis. Next, we showed that telomere length patterns across chromosome ends are highly conserved across tissues and genetic ancestries. Our results extend the blood-based observation of conserved telomere length patterns by Karimian *et al.* (2024) to multiple tissues and ancestries. At the sequence level, we identified distinct repeat signatures associated with different telomere maintenance pathways. TTTGGG repeats are enriched in telomerase-positive cells and serve as strong markers of telomerase activity, whereas ALT-positive cancer cells exhibit complete interspersion of TVRs across all telomeres. Building on earlier work by Lee *et al*. (2014), Sieverling *et al.* (2020), and Conomos *et al.* (2012) on altered TVRs in telomerase- and ALT-positive cells, we show that telomerase activity preferentially introduces *de novo* TTTGGG repeats and that ALT-positive cells have fully interspersed TVRs along all telomeres.

Our tissue dataset included only six individuals, which limits the sensitivity to detect subtle tissue-specific differences in chromosome-specific telomere length patterns. However, the expanding SMaHT cohort will provide increased sample diversity to validate and refine these findings. Similarly, while cancer cell lines offer valuable models for studying ALT-associated telomere dynamics, validating these observations in primary tumor samples will be important for broader relevance.

Despite these constraints, our study highlights the utility of ONT WGS for telomere analysis. We demonstrated that structural variants in genes like *TERT* and *NHP2* associate with genome-wide telomere shortening. Moreover, the consistent chromosome-end-specific telomere length pattern—present despite various age, ancestry, and tissue-specific factors—suggests an underlying conserved maintenance mechanism. For instance, shortening of Xp, which we have identified as typically having the longest telomere, has been linked to breast cancer risk [16], raising the possibility that selectively maintaining such ends could mitigate disease risk. Finally, our findings introduce practical sequence-based markers for telomere biology. TTTGGG repeats provide a quantitative signature of telomerase activity, while the pervasive TVR pattern in ALT cells offers a sensitive, single-molecule indicator for ALT that could be exploited clinically.

## Methods

### Study cohorts and datasets

We analyzed three primary data sets. (1) tissues from deceased human donors collected as part of the SMaHT consortium. Data are accessible through the SMaHT data portal (https://data.smaht.org/) for researchers with approved dbGaP authorization under the study accession number phs004193. (2) ONT sequencing data from the 1000 Genomes Project, with aligned reads and Sniffles2 variant calls downloaded from https://s3.amazonaws.com/1000g-ont/index.html?prefix=ALIGNMENT_AND_ASSEMBLY_DATA/100_PLUS/. (3) telomere-enriched ONT sequencing of fibroblasts, iPSCs, and cancer cell lines generated by Schmidt *et al.* (2024), available via SRA under accession number PRJNA1040425.

### Sample preparation, sequencing, and processing of SMaHT data

The SMaHT tissues utilized in this study were homogenates of liver, lung, colon, and dorsolateral prefrontal cortex from post-mortem donors. Detailed protocols are provided in the SMaHT Network paper [40]. Below is a summary of long-read WGS performed by the different sequencing centers.

#### PacBio HiFi Sequencing

Genomic DNA was generally fragmented to a mode of 18 kb to 22 kb. Libraries were prepared using the SMRTbell Prep Kit 3.0 and subsequently subjected to size selection via the Sage PippinHT instrument to enrich for fragments, typically starting at a minimum of 15 kb (e.g., 15 kb - 17 kb at WashU). Sequencing was conducted on the PacBio Revio platform, targeting a high coverage of 90x to 96x for tissue samples.

#### ONT Sequencing

High molecular weight DNA was extracted, and variable fragmentation targets were used: NYGC aimed for a target size of 45 kb, while BCM targeted 15 kb - 20 kb for tissues. Libraries were prepared using the ONT Ligation Sequencing Kit V14 (SQK-LSK114). The fragmented DNA was also size-selected using methods like the Sage PippinHT (at BCM, using 6 kb - 10 kb or 15 kb - 20 kb High-Pass definitions) or a 3x AMPure XP buffer exchange clean-up (at NYGC). Sequencing was performed using R10.4.1 flow cells on PromethION systems, with a target coverage of approximately 90x at BCM.

### Chromosome-specific telomere length analysis

Telogator2 [26] was used with the recommended settings (-r ont -n 4) for ONT data and (-r hifi -n 4) for PacBio HiFi data, generating per-read telomere lengths for QC-passing telomeric reads and assigning each read to its corresponding chromosome end. Chromosome ends were included only if they contained at least 10 telomeric reads, excluding the acrocentric short arms 13, 14, 15, 21, 22, and Y. Unless otherwise stated, the telomere length reported for a given chromosome end was the mean length across all telomeric reads assigned to that end.

### TVR analysis

A custom computational pipeline starts from BAM files containing telomeric long reads aligned to the T2T-CHM13v2.0 reference genome using minimap2 with parameters -Y -L --eqx --secondary=no -ax map-ont. Optionally, telomere length annotations (e.g., from Telogator2) can be included to facilitate positional mapping of TVRs relative to the telomere start. In the absence of such annotations, reads are filtered based on a minimum number of canonical telomeric repeats. TVR motifs can either be explicitly specified or automatically generated as all possible permutations of triplet repeats followed by ‘GGG’, excluding the canonical ‘TTAGGG’. Two sequence context modes are available: a ‘singleton’ mode, in which TVRs must be flanked by canonical repeats on both sides, and an ‘arbitrary’ mode that allows more flexible flanking contexts. For each read, the nucleotide sequence and base quality scores are extracted. All occurrences of TVRs are identified in both forward and reverse complement orientations. For each detected motif, the base quality scores across the motif region are summed to support downstream filtering based on sequence confidence.

When telomere length annotations are available, the relative position of each TVR within the read is normalized by the annotated telomere length. This normalization allows classification of TVRs into subtelomeric or intratelomeric regions, based on their relative distance from the telomere boundary. Subsequent downstream analysis of the resulting TVR counts was performed using a custom R script. TVRs were filtered based on their context (e.g., ‘arbitrary’), flanking motif length (e.g., three consecutive hexamers), and a minimum summed base quality score (e.g., ≤ 120) to ensure reliability. Based on orientation and relative position, TVRs were classified as subtelomeric, intratelomeric, or artifact. To normalize for differences in sequencing depth and telomere content, TVR counts were expressed per million bps of telomeric or subtelomeric sequence.

### SV analysis

SV calls previously generated for the 1000 Genomes Project ONT cohort [27] were used. In their study, phased SVs were identified using Sniffles2 on haplotagged BAM files. To examine the potential disruption of telomere-related genes, a BED file was generated containing the coordinates of known telomere biology genes, including *TERT, TERC, DKC1, NOP10, NHP2, GAR1, NAF1, PARN, ZCCHC8, TERF1, TERF2, TERF2IP, POT1, TPP1, TINF2, RTEL1, WRN, BLM, CTC1, STN1, and TEN1*, extended by 150 bps upstream of the transcription start site to account for promoter regions. This gene set was intersected with SV breakpoint coordinates to identify potentially gene-disrupting variants.

### Machine learning models for ALT classification

To predict whether individual reads originated from ALT-positive samples, we trained machine learning models using default parameters based on TVRs and telomere length features. For the TVR-based classifier, a random forest model was implemented in R using the caret package, with the relative counts and telomere distance of the TVRs CGAGGG, CTAGGG, GTAGGG, TCAGGG, TGAGGG, TTCGGG, TTGGGG, and TTTGGG per read as input features. For comparison, three models based solely on telomere length were trained using glmnet in R with default settings: one classifying single reads and two aggregating information across 10 or 100 reads by taking the mean telomere length as the predictor.

### Statistical analysis

Statistical analyses were performed in R. Kendall’s tau correlation was used to assess concordance in the rank ordering of telomere lengths across chromosome arms, whereas Pearson correlation was used for comparing absolute agreement between measurements, such as between platforms. Wilcoxon rank-sum tests were applied for independent groups comparisons. An alpha level of 0.05 was used as the threshold for significance, with Benjamini–Hochberg correction for multiple testing. Significance levels were reported using the standard notation: ns (not significant), * (p ≤ 0.05), ** (p ≤ 0.01), *** (p ≤ 0.001), and **** (p ≤ 0.0001).

## Conflict of interest

The authors declare no potential conflicts of interest.

## Code availability

Code for the telomere analysis of long-read WGS data is available from https://github.com/niklas-engel/long-read_telomere_profiling_2025.

## Notes

### Competing Interest Statement

The authors have declared no competing interest.

## References

1. Counter, C. M. et al. Telomere shortening associated with chromosome instability is arrested in immortal cells which express telomerase activity. The EMBO Journal 11, 1921–1929 (1992).

2. Moyzis, R. K. et al. A highly conserved repetitive DNA sequence, (TTAGGG)n, present at the telomeres of human chromosomes. Proceedings of the National Academy of Sciences 85, 6622–6626 (1988).

3. Allshire, R. C., Dempster, M. & Hastie, N. D. Human telomeres contain at least three types of G–rich repeat distributed non-randomly. Nucleic Acids Research 17, 4611–4627 (1989).

4. Harley, C. B., Futcher, A. B. & Greider, C. W. Telomeres shorten during ageing of human fibroblasts. Nature 345, 458–460 (1990).

5. Hayflick, L. & Moorhead, P. S. The serial cultivation of human diploid cell strains. Experimental Cell Research 25, 585–621 (1961).

6. Fagagna, F. d’Adda di et al. A DNA damage checkpoint response in telomere-initiated senescence. Nature 426, 194–198 (2003).

7. Greider, C. W. & Blackburn, E. H. Identification of a specific telomere terminal transferase activity in Tetrahymena extracts. Cell 43, 405–413 (1985).

8. Kim, N. W. et al. Specific association of human telomerase activity with immortal cells and cancer. Science 266, 2011–2015 (1994).

9. Feng, J. et al. The RNA component of human telomerase. Science 269, 1236–1241 (1995).

10. Zijlmans, J. M. J. M. et al. Telomeres in the mouse have large inter-chromosomal variations in the number of T2AG3 repeats. Proceedings of the National Academy of Sciences 94, 7423–7428 (1997).

11. Martens, U. M. et al. Short telomeres on human chromosome 17p. Nat Genet 18, 76–80 (1998).

12. Karimian, K. et al. Human telomere length is chromosome end–specific and conserved across individuals. Science 384, 533–539 (2024).

13. Hemann, M. T., Strong, M. A., Hao, L.-Y. & Greider, C. W. The Shortest Telomere, Not Average Telomere Length, Is Critical for Cell Viability and Chromosome Stability. Cell 107, 67–77 (2001).

14. Xu, Z., Duc, K. D., Holcman, D. & Teixeira, M. T. The Length of the Shortest Telomere as the Major Determinant of the Onset of Replicative Senescence. Genetics 194, 847–857 (2013).

15. Samper, E., Flores, J. M. & Blasco, M. A. Restoration of telomerase activity rescues chromosomal instability and premature aging in Terc−/− mice with short telomeres. EMBO reports 2, 800–807 (2001).

16. Zheng, Y.-L., Zhou, X., Loffredo, C. A., Shields, P. G. & Sun, B. Telomere deficiencies on chromosomes 9p, 15p, 15q and Xp: potential biomarkers for breast cancer risk. Human Molecular Genetics 20, 378–386 (2011).

17. Samassekou, O. et al. Chromosome Arm-Specific Long Telomeres: A New Clonal Event in Primary Chronic Myelogenous Leukemia Cells. Neoplasia 13, 550–IN17 (2011).

18. Ferrer, A., Stephens, Z. D. & Kocher, J.-P. A. Experimental and Computational Approaches to Measure Telomere Length: Recent Advances and Future Directions. Curr Hematol Malig Rep 18, 284–291 (2023).

19. Barthel, F. P. et al. Systematic analysis of telomere length and somatic alterations in 31 cancer types. Nat Genet 49, 349–357 (2017).

20. Li, C. et al. Genome-wide Association Analysis in Humans Links Nucleotide Metabolism to Leukocyte Telomere Length. Am J Hum Genet 106, 389–404 (2020).

21. Taub, M. A. et al. Genetic determinants of telomere length from 109,122 ancestrally diverse whole-genome sequences in TOPMed. Cell Genomics 2, 100084 (2022).

22. Logsdon, G. A., Vollger, M. R. & Eichler, E. E. Long-read human genome sequencing and its applications. Nat Rev Genet 21, 597–614 (2020).

23. Sholes, S. L. et al. Chromosome-specific telomere lengths and the minimal functional telomere revealed by nanopore sequencing. Genome Res. 32, 616–628 (2022).

24. Tham, C.-Y. et al. High-throughput telomere length measurement at nucleotide resolution using the PacBio high fidelity sequencing platform. Nat Commun 14, 281 (2023).

25. Schmidt, T. T. et al. High resolution long-read telomere sequencing reveals dynamic mechanisms in aging and cancer. Nat Commun 15, 5149 (2024).

26. Stephens, Z. & Kocher, J.-P. Characterization of telomere variant repeats using long reads enables allele-specific telomere length estimation. BMC Bioinformatics 25, 194 (2024).

27. Gustafson, J. A. et al. High-coverage nanopore sequencing of samples from the 1000 Genomes Project to build a comprehensive catalog of human genetic variation. Genome Res. 34, 2061–2073 (2024).

28. Coorens, T. H. H. et al. The Somatic Mosaicism across Human Tissues Network. Nature 643, 47–59 (2025).

29. Aubert, G., Baerlocher, G. M., Vulto, I., Poon, S. S. & Lansdorp, P. M. Collapse of Telomere Homeostasis in Hematopoietic Cells Caused by Heterozygous Mutations in Telomerase Genes. PLOS Genetics 8, e1002696 (2012).

30. Tan, K.-T., Slevin, M. K., Meyerson, M. & Li, H. Identifying and correcting repeat-calling errors in nanopore sequencing of telomeres. Genome Biology 23, 180 (2022).

31. Ding, Z. et al. Estimating telomere length from whole genome sequence data. Nucleic Acids Research 42, e75 (2014).

32. Lee, M. et al. Telomere extension by telomerase and ALT generates variant repeats by mechanistically distinct processes. Nucleic Acids Research 42, 1733–1746 (2014).

33. Bryan, T. M., Englezou, A., Dalla-Pozza, L., Dunham, M. A. & Reddel, R. R. Evidence for an alternative mechanism for maintaining telomere length in human tumors and tumor-derived cell lines. Nat Med 3, 1271–1274 (1997).

34. Heaphy, C. M. et al. Prevalence of the Alternative Lengthening of Telomeres Telomere Maintenance Mechanism in Human Cancer Subtypes. The American Journal of Pathology 179, 1608–1615 (2011).

35. Cesare, A. J. & Reddel, R. R. Alternative lengthening of telomeres: models, mechanisms and implications. Nat Rev Genet 11, 319–330 (2010).

36. Akincilar, S. C., Unal, B. & Tergaonkar, V. Reactivation of telomerase in cancer. Cell Mol Life Sci 73, 1659–1670 (2016).

37. Conomos, D. et al. Variant repeats are interspersed throughout the telomeres and recruit nuclear receptors in ALT cells. Journal of Cell Biology 199, 893–906 (2012).

38. Sieverling, L. et al. Genomic footprints of activated telomere maintenance mechanisms in cancer. Nat Commun 11, 733 (2020).

39. Zhang, J.-M. & Zou, L. Alternative lengthening of telomeres: from molecular mechanisms to therapeutic outlooks. Cell Biosci 10, 30 (2020).

40. The Somatic Mosaicism across Human Tissues Network. Comprehensive benchmarking of somatic mutation detection by the SMaHT Network. 2025.10.09.678885 Preprint at 10.1101/2025.10.09.678885 (2025).

