## Supplementary Information for "Nanopore whole-genome sequencing reveals conserved chromosome-specific telomere architecture across tissues and populations"

### Supplementary Figures

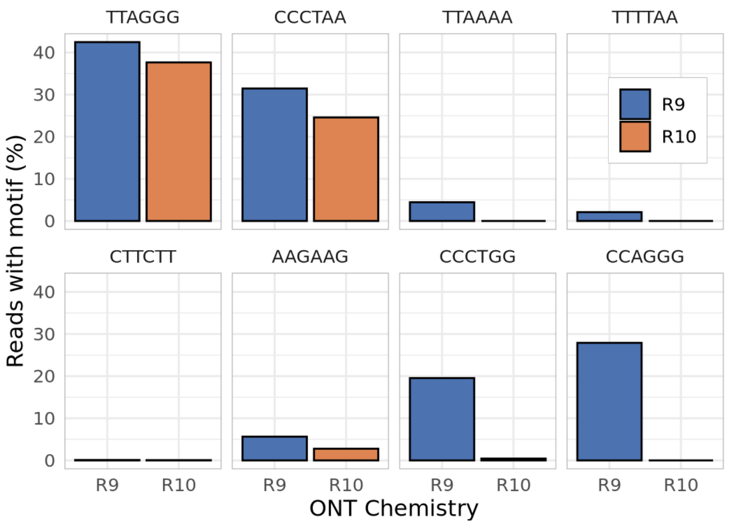


***Supplementary Figure 1. Basecalling error motifs in telomeric reads across Nanopore chemistries.***
*Barplot of the proportion of ONT telomeric reads containing either the canonical TTAGGG motif or known basecalling error motifs (TTAAAA, CTTCTT, CCCTGG), stratified by R9 and R10 ONT chemistries.*

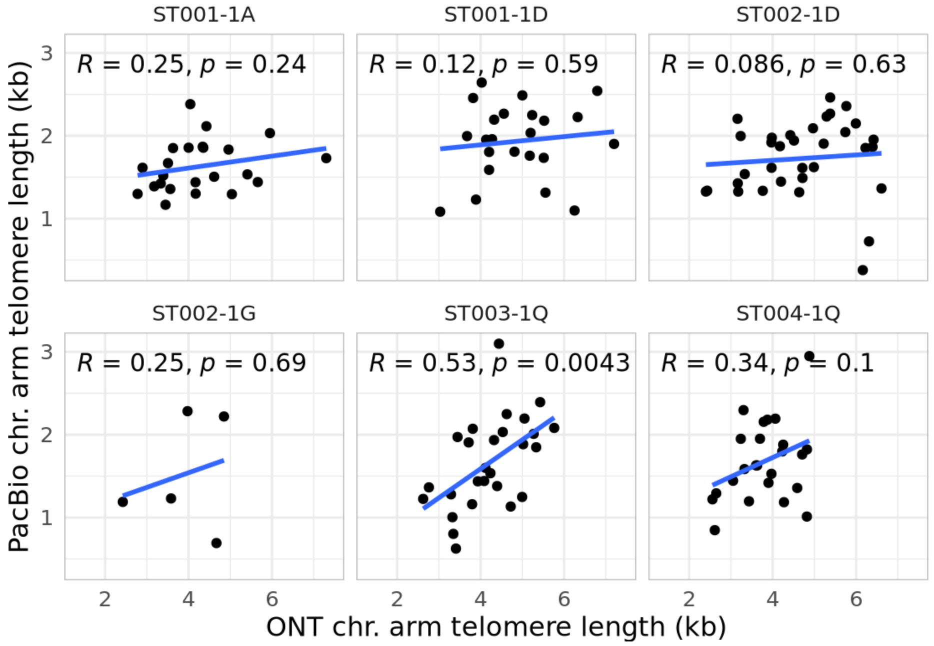


***Supplementary Figure 2. Comparison of telomere length estimates from ONT and PacBio HiFi data.***
*Correlation of chromosome-specific telomere length estimates between samples sequenced with both ONT and HiFi at the same sequencing center. Blue lines indicate linear regression fits; Pearson correlation coefficients and p-values are shown in each panel.*

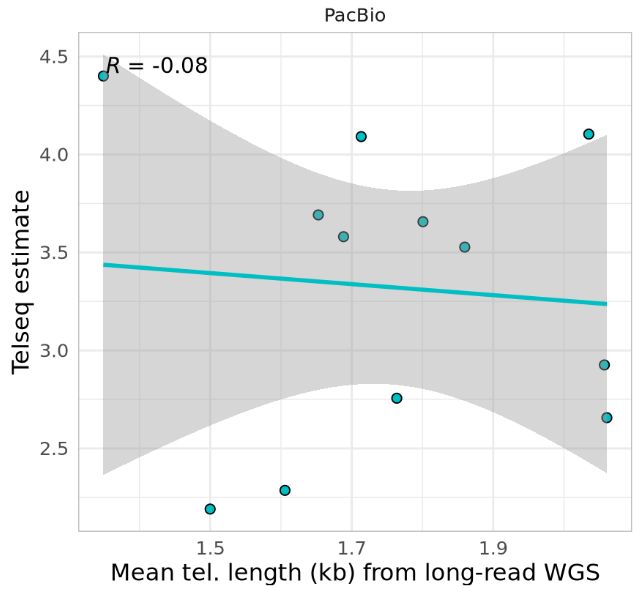


***Supplementary Figure 3. Correlation of telomere length estimates between PacBio and short-read WGS.***
*Correlation of mean telomere length estimates from PacBio HiFi with matched short-read WGS estimates using Telseq. Regression lines and Pearson correlation coefficients are shown.*

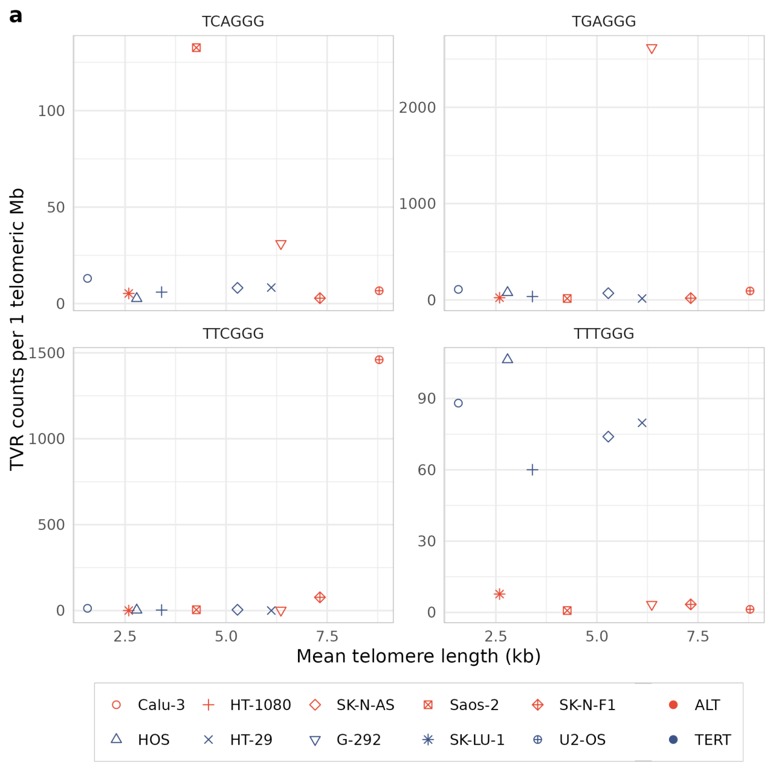


***Supplementary Figure 4. TVRs in ALT+ and TERT+ cell lines.***
*Number of singleton TVRs per 1 Mb of telomeric sequence for TCAGGG, TGAGGG, TTCGGG, and TTTGGG repeats, plotted against the mean telomere length (kb) for each sample. ALT+ and TERT+ status is indicated by color; cell lines are distinguished by shape.*
